# Not-so-great tit: early-life environment drives long-term decrease in adult body mass in a wild bird population

**DOI:** 10.64898/2026.02.11.705378

**Authors:** David López-Idiáquez, Ella F. Cole, Devi Satarkar, Samuel J. Crofts, Keith McMahon, Ben C. Sheldon

## Abstract

Body mass is a key organismal characteristic that impacts many physiological and ecological processes and often a strong determinant of fitness. Recent studies have documented temporal phenotypic changes in this trait in many populations, but identifying the mechanisms underpinning these changes can be difficult. Here, we use 47 years of data to analyse how adult and nestling body mass have changed over time in a great tit *Parus major* population in Wytham Woods (UK). Further, we link those changes to three environmental variables previously recognised as drivers of body mass: temperature, intra- and inter-specific competition and temporal mismatch with a key prey during breeding, winter moth Operophtera brumata caterpillars. Temporal analyses of adult body mass revealed contrasting dynamics at the between- and within-cohort levels, mirroring Simpson’s Paradox. At the population level we report a marked decrease in body mass in adults between 1978 and 2024 (−0.042 Haldanes), and show that this results from phenotypic plasticity, driven by a negative between-cohort trend likely reflecting carry-over effects of the early environment. Within cohorts, however, trends were consistently positive likely reflecting an age-dependent mass increase. The change in adults was paralleled by a change in nestling body mass (−0.036 Haldanes). Nestling mass was negatively associated with estimated intensity of intraspecific competition, as well as inter-specific competition from blue tits *Cyanistes caeruleus*, as quantified by local population density. These effects carried over to adulthood, as shown by a negative association between adult mass and the population density experienced at early life. Temperature during development and mismatch with the caterpillar food supply, despite being associated with adult and nestling mass, did not explain the observed declines in mass, largely because these have not changed over time. Overall, our results illustrate the potential for effects mediated early in development to carry-over into long-term phenotypic change at later life history stages, and emphasise the value of considering multiple effects as drivers of phenotypic change in natural populations.

## Introduction

Phenotypic responses to environmental change are ubiquitous across taxa. One of the main environmental drivers of these changes is rapid global warming, and many studies have reported phenotypic and genetic responses in traits such as morphology (Sheridan & Bickford, 2011), ornamentation (López-Idiáquez et al., 2022), and phenology (Thackeray et al., 2016). In addition, other abiotic and biotic factors, such as resource availability (Arcese & Smith, 1988), predation intensity (Gosler et al., 1995), or extreme climatic events (Satarkar et al. 2025), also play important roles as drivers of phenotypic change. Analysing how phenotypes respond to environmental change and the drivers and mechanism behind those responses is, therefore, a crucial step in understanding how populations will cope with these shifts. For instance, while short-term adaptation to fluctuating environmental conditions can be achieved through phenotypic plasticity, only microevolutionary changes of a trait or its plasticity can ensure the adaptation to continued directional environmental variation (Oostra et al., 2018; Van Buskirk & Steiner, 2009).

In this context, body mass is of particular interest due to its relevance as an organismal trait, with studies both across- and within-species, showing that it can predict the expression of fitness-related traits such as survival, fecundity, or metabolism (Dumas et al., 2024; Maldonado-Chaparro et al., 2015). Although body mass is tightly linked to structural size and has a moderate to strong genetic basis (Araya-Ajoy et al., 2019; Dumas et al., 2024; Garant et al., 2004), it is also an inherently plastic trait sensitive to variation in the environmental conditions (Macleod et al., 2005; Moiron et al., 2018). Therefore, body mass responses to environmental change may be the product of both phenotypic plasticity and a genetic change (Gienapp et al., 2008).

Most research conducted to date analysing long-term datasets on body mass or size has focused on testing a prediction derived from an intraspecific extension of Bergmann’s rule – i.e., that decreasing body sizes reflect a response to increasing temperatures (He et al., 2023; Teplitsky & Millien, 2014). Results from these studies generally conform with Bergmann’s rule, as they report declines in body size or mass in association with temperature increase (Sheridan & Bickford, 2011; Youngflesh et al., 2022). For instance, warming has been proposed to drive a general decrease in body mass of 105 bird species across North America (Youngflesh et al., 2022). The rationale behind this temperature-mass association is that smaller, or lighter-bodied, individuals have lower cooling costs than their larger or heavier counterparts (Gardner et al., 2011; Youngflesh et al., 2022). However, this view has recently been challenged, as the shifts in mass and size shown by most bird populations are generally too small to impact heat production and dissipation (Nord et al., 2024). This implies that the effects of temperature on body mass may be weaker than previously thought, or that they might reflect variation of non-measured environmental variables (Husby et al., 2011; Wilcox et al., 2024). For instance, in three Dutch great tit *Parus major* populations, the decrease in adult mass between 1979 and 2008 was not linked to changes in temperature, despite the warming in the area, but rather to an increasing temporal mismatch with their main prey during breeding (Husby et al., 2011). Furthermore, very few studies have found adaptive changes in body size as a response to warming, as body size is responding to environmental change mostly through phenotypic plasticity rather than microevolution (Teplitsky & Millien, 2014). This calls for more research to clarify the role of ambient temperature change as a driver of temporal trends in body mass and underscores the need of considering alternative environmental factors in addition to temperature.

Besides the potential physiological advantages of declines in body size as a response to warming, a complementary factor explaining trends in body mass may be changes in resource availability, as is well established that higher resource availability is linked to higher body mass and condition (Cox et al., 2019; Husby et al., 2011; Wilcox et al., 2024). The effects of resource availability can also arise indirectly and factors such as intra- and inter-specific competition can limit energy availability by competitive process with neighbours (Arcese & Smith, 1988). This was experimentally demonstrated in a song sparrow *Melospiza melodia* population, where supplementary feeding of a subset individuals buffered the reduction in reproductive success, compared to control individuals, caused by a sixfold increase of population density (Arcese & Smith, 1988). Therefore, we can expect a link between body mass and population density, an association reported in several taxa (Brown et al., 2022; Garant et al., 2004; Pettorelli et al., 2002). Two of the main mechanisms that can explain the reduced energy availability at higher densities and its effects on body mass are density effects on foraging behaviour (Cresswell, 1998; McMahon & Marples, 2017) and parasite transmission (Albery et al., 2024). For instance, blackbirds *Turdus merula* experience a reduction in foraging efficiency when conspecific numbers in the foraging patch are higher (Cresswell, 1998), and parasite prevalence in juvenile Soay sheep *Ovis aries* is positively correlated with population density (Albery et al., 2024). Regardless of the mechanism, these patterns highlight the relevance of considering intra- and inter-specific competition, alongside other ecological drivers, when analysing body mass responses to environmental variation.

Phenotypic responses to changes in environmental conditions do not only occur in the short term; they can also operate over longer periods of time, carrying over from one life-stage to another. In this context, the effects of the environmental conditions experienced at early-life are of particular relevance (Monaghan, 2008). There is ample experimental and correlative evidence showing that the environmental conditions during early life can impact adult performance, influencing traits such as life-span (Reid et al., 2003), ageing (Spagopoulou et al., 2020), and reproductive success (Lindström, 1999; van De Pol et al., 2006). For instance, chough *Pyrrhocorax pyrrhocorax* fledglings that develop under favourable environmental conditions have longer lifespans, allowing them to achieve higher lifetime reproductive success compared to individuals raised in harsher environments (Reid et al., 2003). These long-lasting effects of the environment experienced at early life reveal the potential complexity of underpinning the mechanisms behind the trends in phenotypic traits we see in nature and highlight the relevance of analysing the effects of different environmental variables at different life-stages. For instance, previous studies in birds and other species have shown that early-life effects can carry over into adulthood to affect body mass (Brown et al., 2022; Pettorelli et al., 2002) and that, in some cases, such effects can account for up to 40% of the variation in adult mass (Pettorelli et al., 2002). Studies analysing temporal trends in adult body mass, however, focus mostly on the environmental effects experienced during adulthood, missing a potentially relevant piece for our understanding of the dynamics of this trait.

In this study, we use more than 17,700 measures from wild great tits to analyse the temporal trends in adult body mass during breeding over almost five decades. We analyse how body mass has changed between 1978 and 2024 at both the phenotypic and genetic levels using a social pedigree, and link those changes to variation in temperature, and intra- and inter-specific competition with blue tits - a confamilial species with an overlapping ecological niche (Perrins, 1979). In addition, we leverage 38 years of data on the mismatch between great tit timing of breeding and a key food resource, winter moth *Operophtera brumata* caterpillars, to explore the links between food availability and mass in adults and nestlings. To analyse the role of early-life conditions in shaping adult body mass, we analyse trends in 77,787 individual observations of nestling mass, how they are influenced by the abovementioned environmental variables, and their association with adult mass. Because both the number of breeding great and blue tits and temperature have increased in our study area, we predict a reduction in adult body mass during the study period, regardless of whether these drivers act directly on adults or indirectly through early-life effects. Finally, based on previous evidence in other systems (Teplitsky & Millien, 2014) we predict that most of the change in adult body mass will occur at the phenotypic and rather than at the genetic level.

## Methods

### Study system

We studied the great tit population in Wytham Woods, a mixed forest near Oxford that has been continuously monitored since 1947, and where, since 1960, approximately 1000 nest boxes have been available for great tits to breed (Perrins, 1965). Wytham is divided into nine sections (hereafter: section) which, although subjectively delimited, still capture environmental differences between them (Minot & Perrins, 1986). During the breeding season (April-June), nest boxes are visited at least once a week to collect detailed phenological, breeding and morphological data. Adults are captured in the nest boxes when nestlings are between 10 and 14 days old. At capture, every adult is measured to record body mass (to the nearest 0.1 g) and aged based on plumage characteristics (Svensson, 1992). Every bird is marked with an individually numbered metal ring and, from 2011 onwards, with a RFID tag. These RFID tags allow us to identify the adults without physically capturing them, and consequently, since 2011 onwards, adult capture effort was reduced and focused on untagged individuals. We analysed the potential impacts of this sampling approach and found no evidence that it biased the trends reported below (see supplementary material (SM)-1). Nestlings are ringed and weighted when they are 14 days old, time at which their growth and mass typically plateaus (van Balen, 2002). Adult body mass shows a positive association with the mass the individual had as a nestling (0.387±0.010, F_1, 6101.7_=1301.7, p<0.001, n=9306). In the Wytham Woods great tit population a generation time has been estimated as 1.81 years (Bouwhuis et al., 2009). In this study we used data from 1978 onwards, because by the end of the 1970s the effects of sparrowhawks *Accipiter nisus* returning to Wytham on great tit mass had stabilised (see Gosler et al. 1995 for further details).

Data on winter moth half-fall dates were provided by Dr L. Cole. Half-fall date is the day on which 50% of the seasonal total winter moth caterpillars are collected in water traps under oak trees, and provides a robust index of the timing of peak of food availability for great tits (Hinks et al., 2015; Morley et al., 2025; Wilkin et al., 2009).

### Statistical analysis

#### Temporal trends in body mass

We analysed temporal trends in adult and nestling great tit body mass using linear mixed models (LMM). For adults, we fitted one LMM in which body mass was included as dependent variable and year of measurement (hereafter year; as a continuous variable), sex, age (1-year-old vs >1-year-old) and section as fixed effects. Year (as a categorical variable) and individual identity were included as random effects. We complemented this analysis by fitting a Bayesian random slope model. This model included mass as a fixed effect, and year (as a continuous variable), sex and section as fixed effects. Besides the model also included year (as a categorical variable) and individual identity as random effects and a random slope between year (as a continuous variable) and year of birth (as a categorical variable) to test for cohort-specific temporal patterns. We used weakly informative priors, for the fixed effects we used Normal (0, 2) priors and a Normal (18, 5) prior for the intercept. For random effects we used Normal (0, 1) priors and an LKJ (2) prior for the correlation between the intercept and random slope.

We analysed temporal trends in nestling body mass by fitting two LMMs. First, we analysed temporal trends in nestling mass using all nestlings; this model included nestling mass as a dependent variable and year (as a continuous variable) and section as fixed effects. As random effects, we included year (as a categorical variable), and nest box and brood identity. Second, we analysed temporal trends in the subset of the nestlings that recruited to the population as breeders, to explore whether viability selection on nestling mass might have also changed in our study period. This model had the same structure as the previous one but included only those birds identified as breeding adults in the population in subsequent years. To identify those nestlings recruiting from the 2024 cohort, we used the breeding data from 2025.

#### Strength of selection on nestling mass

To formally test for changes in selection strength on nestling mass, we computed yearly standardised selection differentials by subtracting the average standardised mass of the nestlings in a certain year to the average standardised mass of those that recruit to the population as breeders. We then regressed those values against year in a linear model to analyse the temporal trends in the selection differentials on body mass.

#### Linking body mass and temperature

We analysed the link between temperature and body mass using two complementary approaches. First, we linked adult mass at capture (t) to the average spring (April_t_ to June_t_) and previous winter (October_t-1_ to January_t_) temperature in separate models. For adults, the LMMs included body mass as the dependent variable, and spring or winter temperature, year (as a continuous variable), section, sex and age (1-year-old vs. >1-year-old) as explanatory terms. Year (as a categorical variable) and individual identity were included as random effects. For nestlings, the LMMs included body mass of all nestlings as the dependent variable, and spring or previous winter temperature, year (as a continuous variable) and section as explanatory terms, with year (as a categorical variable) and nest and brood identity being included as random effects.

Second, to better capture any short-term effects of temperature on body mass, we used average temperature over a relative rather than a fixed interval. For adults, this window represented the average temperature experienced by each individual 10 days prior to capture. For nestlings, it corresponded to the average temperature experienced by each brood between hatching and 15 days post-hatching (i.e., the general ringing date for most broods). Model structure for both adults and nestlings was the same as in the fixed-interval analysis, except that temperature effects on body mass were modelled using natural cubic splines with five degrees of freedom, rather than as a linear term, to capture potential non-linear associations between body mass and temperature, which were not expected in the previous section.

We obtained daily temperature data between 1978 and 2024 from the Central England dataset (Met Office Hadley Centre - www.metoffice.gov.uk/hadobs/).

#### Linking body mass and intra- and inter-specific competition

We calculated intra- and inter-specific competition as the number of great and blue tit breeding pairs around each nest. We considered blue tits in addition to great tits as these two species notably overlap in their ecological niches, and although great tits are expected to dominate over blue tits, blue tits can still impose a pressure on great tits (Gamelon et al., 2019; Minot, 1981). We computed breeding density in 17 buffers, from 100 meters to 4000 meters around each nest (i.e. from small-scale territory level effects to population density), as competition effects can differ depending on the considered spatial scale. With this information we fitted one LMM per buffer and species to explore the effects of great and blue tit numbers on body mass. For adults, the models included body mass as dependent variable and the number of great tits or blue tits at each buffer, year (as a continuous variable), section, sex and age (1 year-old vs. >1 year-old) as explanatory variables. Here, we included year (as a categorical variable) and individual identity as random effects. In nestlings, we included body mass of all nestlings as explanatory variable and the number of great tits or blue tits at each buffer and year (as a continuous variable) and section as dependent variables. These models included year (as a categorical variable), nestbox, and brood identity as random effects.

#### Linking body mass and mismatch

We computed mismatch between the peak of food availability and the peak of food demand as the number of days between the date when nestlings are 10 days old (i.e. peak of food demand) and winter-moth caterpillar half-fall date. We then analysed the association of mismatch with adult and nestling body mass using LMM. For both adult and nestling models, body mass was included as the dependent variable and the effects of mismatch were modelled by fitting a natural cubic spline with five degrees of freedom. In the adult body mass model, year (as a continuous variable), sex and section were included as covariates and year (as a character) and individual identity as random factors. In the model for nestlings, we included year (as a continuous variable) and section as covariates and brood identity, nest and year (as a character) as random factors. Winter moth half-fall data was available for 38 out of the 47 years with body mass information (non-available years 1978-1982 and 1989-1992).

#### Temporal trends in the environmental variables

To compute temporal trends in the environmental variables we fitted a series of linear models that included the yearly average environmental value as dependent variable and year (as a continuous variable) as explanatory term.

#### Quantitative genetics of body mass

The long-term monitoring of our population allowed us to construct a social pedigree describing the ancestry of the great tits. This social pedigree assumes that adults identified at the nest are the biological parents of the offspring. Although great tits are socially monogamous, extrapair copulations occur in the species. The rate of extrapair paternity in our population (∼12%, Patrick et al. 2012), however, is not expected to bias the quantitative genetic estimates obtained from analyses of the social pedigree (Firth et al., 2015), which has been successfully used in recent studies in our population (Jones et al., 2025; Satarkar, Sepil, et al., 2025). The pruned version of the pedigree analysed here included 12,762 individuals and extended over 38 generations.

We used this pedigree in an animal model which allowed us to partition the variance of adult body mass (V_P_) into its additive genetic (V_A_), permanent environmental (V_PE_), year (V_YR_) and residual (V_R_) components (Kruuk, 2004). With that aim, we fitted a Bayesian LMM that included adult body mass as the dependent variable and age (1-year-old vs. >1-year-old) and sex as fixed effects. The random structure of the model included: the individual identity linked to the pedigree to estimate V_A_; the individual identity to estimate V_PE_, and year to estimate V_YR_. Narrow-sense heritability was estimated as h^2^=V_A_/V_P_. The model was fitted using default priors.

To estimate the temporal trends in breeding values, which would indicate a change in the genetic component of body mass, we fitted a linear regression of the Best Linear Unbiased Predictors (BLUPs) of V_A_ against the mean breeding year of each individual (Bonnet et al., 2019). We accounted for uncertainty in BLUP estimates by using the full posterior distribution of the model (Hadfield et al., 2010). Specifically, we ran a regression between the BLUP values against the individual mean values of the breeding year for each posterior sample of the model.

In all models the continuous explanatory variables were scaled to a mean of zero and standard deviation of unity. All models have been run in R (R Core Team 2023, v. 4.5.1) using *lme4* (Bates et al. 2015, v. 1.1-37) and *brms* (Bürkner 2017, v. 2.22.0) packages. Bayesian models were run for 10000 interactions across four chains with a warm-up of 1000 iterations and a thin of 10. These values ensured model convergence and good effective sample sizes, which were assessed by inspection of the trace plots and the Gelman-Rubin convergence diagnostic. We calculated predicted trends from the models using *ggeffects* (Lüdecke, 2018) To prune the pedigree and compute its statistics we used the packages *MCMCglmm* (Hadfield 2010, v. 2.36) and *pedtricks* (Martin et al. 2024, v. 0.4.2).

## Results

### Temporal trends in adult and nestling mass

Adult great tit body mass in Wytham Woods has decreased by ∼1 gram between 1978 and 2024, representing a 1.02 standard deviation change (4.96% decrease) in 47 years, which corresponds to a rate of −0.042 Haldanes per generation (Table 1 & Fig. 1A). A more nuanced analysis of the temporal trends, fitting a random slope model, revealed a two-level process by which the population level negative trend pattern masked a positive temporal effect within each great tit cohort (Fig. 1C; see SM-2 for full model results). Nestling body mass has also decreased over the study period, both when considering all birds (−1.4 grams [0.85 std; −7.37% decrease; −0.036 Haldanes per generation]; Table & Fig. 1) and only those that recruited as breeders (−0.94 grams [0.74 std; 4.89% decrease; −0.024 Haldanes per generation]; Table & Fig. 1). Post-hoc analyses revealed a significant difference between the slopes estimated from adults and all nestlings (Wald z-test: z=2.231, p=0.025) but not between the slopes estimated from adults and nestlings that recruited as breeders (z=0.310, p=0.756).

**Table 1:** Results from the linear mixed model analysing temporal trends in adult and nestling great tit body mass. Year was scaled to have a mean of zero and standard deviation of one. Bold denotes statistical significance.

|  | Estimate | SE | F | P |
| --- | --- | --- | --- | --- |
| Adult body mass (n=17785) |  |  |  |  |
| **Year** | **-0.231** | **0.020** | **F<sub>1,50.6</sub>=125.5** | **<0.001** |
| **Sex<sub>male</sub>** | **0.358** | **0.015** | **F<sub>1,11604.5</sub>=557.35** | **<0.001** |
| **Age<sub>juv</sub>** | **-0.256** | **0.011** | **F<sub>1,12555.5</sub>=520.5** | **<0.001** |
| **Section** |  |  | **F<sub>8,13310.3</sub>=10.87** | **<0.001** |
| All nestling body mass (n=77787) |  |  |  |  |
| **Year** | **-0.382** | **0.064** | **F<sub>1,45.40</sub>=35.13** | **<0.001** |
| **Section** |  |  | **F<sub>8,817.51</sub>=31.88** | **<0.001** |
| Recruiting nestling body mass (n=7129) |  |  |  |  |
| **Year** | **-0.246** | **0.043** | **F<sub>1,47.34</sub>=31.63** | **<0.001** |
| **Section** |  |  | **F<sub>8,731.1</sub>=10.76** | **<0.001** |

**Figure 1.**
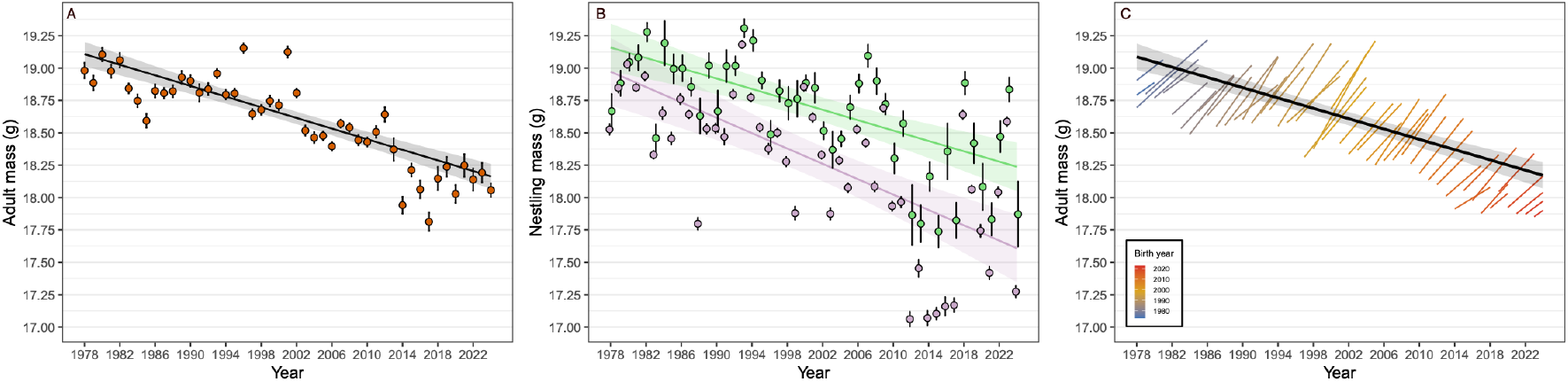
Temporal trends in body mass in breeding adult (A & C) and nestling (B) great tits in Wytham Woods between 1978-2024. In A and B lines and ribbons represent the trends predicted by the model. Dots and whiskers represent the mean ± standard error raw body mass. In B, purple represents the trend of all fledged nestlings and green the trend of those that recruited to the population as breeders. In C, the black line represents the trend in the cohort weighted average (i.e. population level effect – analogous to A) and each line represents the within-cohort temporal trend. Year corresponds to the year in which birds were measured.

### Strength of selection on nestling mass

We found positive selection on nestling mass in all years, with an average selection differential of 0.248 [range: 0.013 – 0.604] and that the strength of selection on nestling mass has increased over our study period (effect of year 0.048±0.020, F1,45=5.879, p=0.019, SM-3).

### Quantitative genetics of body mass

We found that great tit body mass was moderately heritable (h^2^=0.374 [0.334, 0.404]). Consistent environmental differences across individuals (V_PE_), and across years (V_YR_) explained a moderate proportion of the variance in great tit body mass, respectively 0.131 [0.106, 0.162] and 0.114 [0.078, 0.169]. We did not find any evidence of a change in the breeding values of body mass during the study period (effect of year: −0.010 [-0.035, 0.004]; see SM-4 Fig. 1).

### Linking temperature and body mass

Our models analysing the association between seasonal temperature and adult body mass showed that neither spring (F_1,45.5_=0.317, p=0.575) nor winter (F_1, 43.3_=0.001, p=0.994) temperature were significantly associated with adult body mass (see SM-3 for full model results). For nestling body mass, we found a significant and positive association with spring temperature (0.185±0.071, F_1, 44.10_=6.727, p=0.012, see SM-5 for full model results).

When considering temperature relative to adult capture date and nestling development, we found significant non-linear associations with body mass. For adults, there was a significant non-linear association between mean temperature and mass (Table 2), which was negative across the temperature range with some variation in strength (Fig. 2). For nestlings, the model showed a significant non-linear association with temperature, with nestling mass being higher at intermediate temperatures (Table 2, Fig. 2). Temporal trends of both adult and nestling mass remain mostly unchanged after including temperature at capture and development respectively (Tables 1 & 2).

**Table 2:** Results from the linear mixed model including natural cubic splines (df=5) analysing the association between adult body and nestling mass and mean temperature 10 days before capture. Year was scaled to have a mean of zero and standard deviation of one. Bold denotes statistical significance.

|  | Estimate | SE | F | P |
| --- | --- | --- | --- | --- |
| Adult body mass (n=17785) |  |  |  |  |
| Temperature at capture |  |  | **$F_{5,11382.6}=30.81$** | **<0.001** |
| Section |  |  | **$F_{8,13345.3}=10.46$** | **<0.001** |
| Year | -0.236 | 0.020 | **$F_{1,52.4}=129.08$** | **<0.001** |
| Sex<sub>male</sub> | -0.356 | 0.015 | **$F_{1,11614.7}=557.86$** | **<0.001** |
| Age<sub>juv</sub> | -0.251 | 0.011 | **$F_{1,12524.4}=506.51$** | **<0.001** |
| Nestling body mass (n=77787) |  |  |  |  |
| Temp. development |  |  | **$F_{5,9555.0}=22.320$** | **<0.001** |
| Section |  |  | **$F_{8,817.5}=32.285$** | **<0.001** |
| Year | -0.391 | 0.066 | **$F_{1,45.4}=34.590$** | **<0.001** |

**Figure 2.**
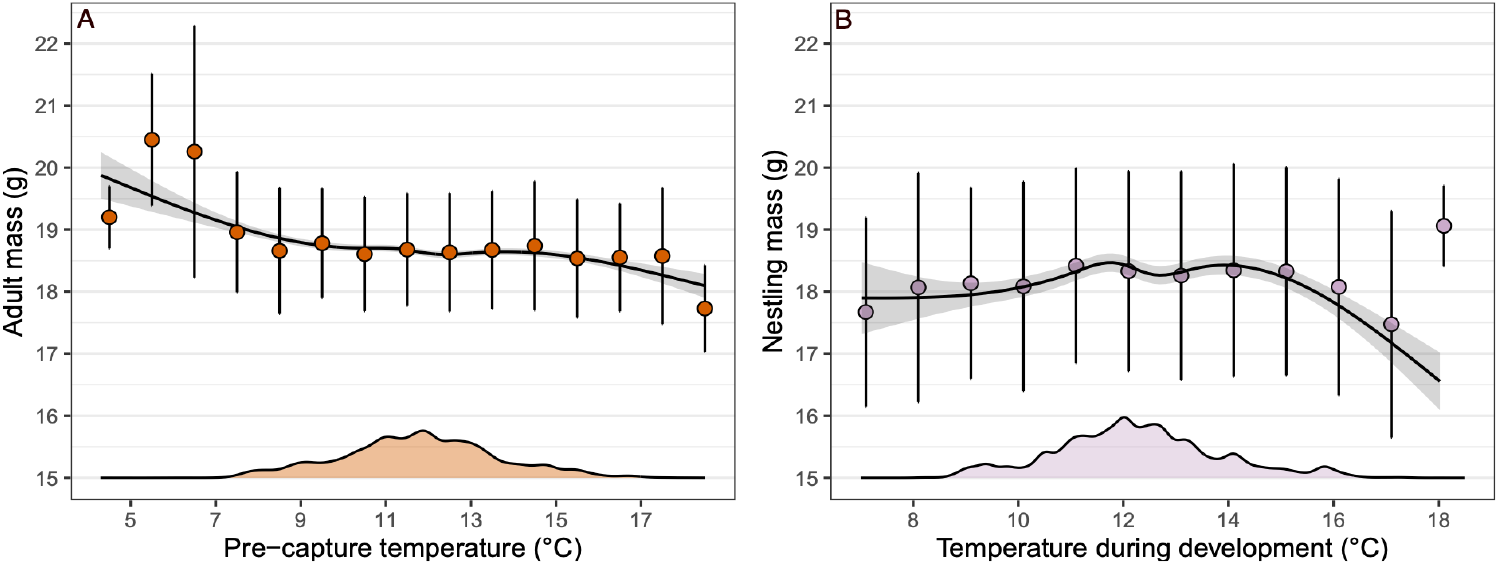
Association between temperature and adult (A) and nestling (B) body mass for great tits in Wytham Woods between 1978-2024. Lines and ribbons represent the splines predicted by the model. Dots and whiskers represent the mean ± one standard deviation raw body mass in 1º C bins. Note that the sample size in each bin differs. Density plots represent the distribution of temperature in each analysis (and hence give information about the sample size). In adults, temperature represents the average temperature 10 days before capture and in nestlings the average temperature from hatching to 15 days after hatching.

### Linking competition and body mass

Analyses of the association between intra- and inter-specific competition and adult body mass at different spatial scales revealed a weak effect of great and blue tit numbers on adult body mass (Fig. 3A, see SM-6 for full model results). Great tit mass showed a positive and significant association with the number of great tits breeding at large spatial scales (3000 and 4000 m). Great tit mass showed a significant and negative association with the number of blue tits at small and intermediate spatial scales (from 100 to 1000 meters), which shifted to being positive in the 2500 meters buffer.

**Figure 3.**
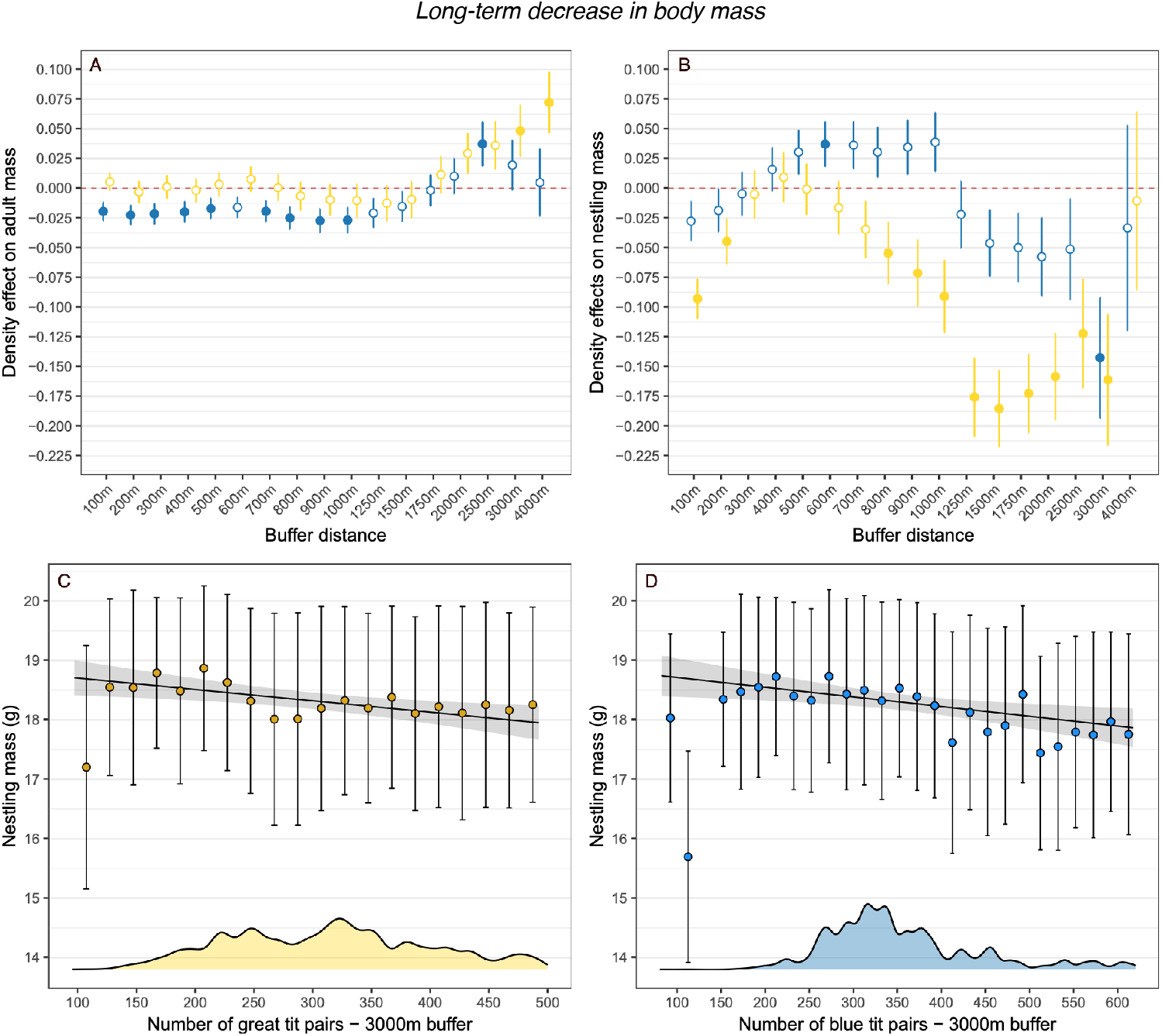
Associations between adult (A) and nestling (B) mass and the number of great (yellow) and blue tit (blue) breeding in buffers of different distances from the focal nest, for great tits in Wytham Woods between 1978-2024. Dots and whiskers represent the estimate of the association between mass and density (scaled) and its standard error. Closed circles represent significant (p<0.05) and open circles non-significant associations. Associations between great tit nestling body mass and the number of great (C) and blue tits (D) breeding in a buffer of 3000 meters around the focal nest. Lines and ribbons represent the trends predicted by the model. Dots and whiskers represent the mean ± one standard deviation raw body mass in bins of 20 pairs. Note that the sample size in each bin differs.

When considering the effects of competition on nestling body mass, number of great tits showed a general negative effect on body mass (Fig. 3B), being strongest at medium and large spatial scales (3000-meter buffer: −0.161±0.055, F_1,180.44_=8.536, p=0.003, Fig. 3C). Blue tit density effects were mostly non-significant across spatial scales (Fig. 3B), except for a weak positive association for the 600-meter buffer and a stronger negative association at 3000 meters (− 0.142±0.050, F_1, 419.43_=7.876, p=0.005, Fig. 3D). For further details on model outputs see SM-6.

Finally, to explore the associations of the early-life environment and adult mass we linked the total number of great and blue tits at 3000 meters experienced as a nestling and the mass of that nestling as an adult, finding a significant and negative association (−0.034±0.015, F_1, 4246.2_=4.88, p=0.027, n=9306, see SM-7 for full model results).

### Linking body mass and mismatch

For adults, we found a negative association between mismatch between the hatching date of the brood they cared for and caterpillar peak date and body mass, adults being heavier when their broods hatched earlier than the peak of food availability. However, this effect was weak and mostly driven by early breeding individuals (Fig. 4A). For nestlings the highest mass was achieved in those broods having a mismatch value close to zero, mostly decreasing at both sides of the peak (Fig. 4B).

**Figure 4.**
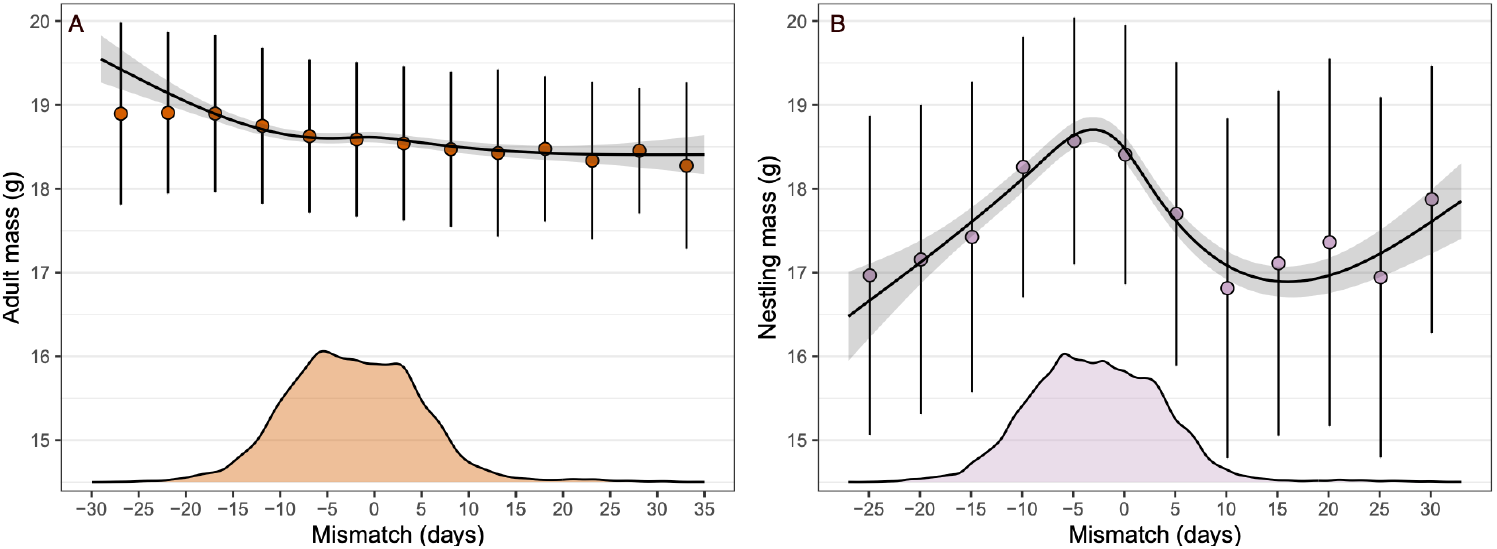
Association between adult (A) and nestling (B) body mass and their mismatch with the peak of food availability for great tits in Wytham woods between 1978-2024. Lines and ribbons represent the splines predicted by the model. Dots and whiskers represent the mean ± one standard deviation of raw body mass in 5-day bins. Note that the sample size of each bin differs. Density plots represent the distribution of mismatch (and hence give information about the sample size) in each analysis.

### Temporal trends in the ecological variables

For temperature, seasonal temperature significantly increased across our study period, both when considering spring (0.034±0.006° C year^-1^, F_1,45_=24.41, p<0.001) and winter (0.033±0.007° C year^-1^, F_1,45_=18.33, p<0.001) temperature. In contrast, yearly average temperatures 10 days before adult capture (−0.006 ±0.014 ° C year^-1^, F_1, 45_=0.242, p=0.625) and nestling development (−0.010± 0.013°C year^-1^, F_1, 45_=0.575, p=0.452) have not shown a significant temporal trend over the study period. The average number of great and blue tits in the 3000-meter buffer has significantly increased during the study period (great tits: 2.175±0.814 pairs year^-1^, F_1, 45_=7.126, p=0.010; blue tits: 4.191±0.714 pairs year^-1^, F_1, 45_=34.38, p<0.001). Finally, mismatch with the peak of food availability has remained stable across the study period (0.037±0.061 days year^-1^, F_1, 36_=0.369, p<0.547).

## Discussion

Using an individually-based dataset comprising 17,785 observations, we show that great tit body mass at breeding has decreased by approximately 1 gram (of the order of 1 s.d.) between 1978 and 2024, a change that can be attributed to phenotypic plasticity. Moreover, the analysis of 77,787 nestling body mass observations revealed a stronger change in mass at early life over the same period. Our results suggest that the decrease in adult body mass is driven by a negative effect of intra- and inter-specific competition on mass at early life, which carries over into adulthood, something supported by our results showing a negative temporal trend in mass between-but not within-cohorts. Body mass of both adult and nestlings was also sensitive to the variation in mismatch with a key prey during breeding and to temperature during breeding. However, these two factors have remained stable during the study period suggesting that they do not explain the reported temporal trends in mass. These findings align with previous work highlighting the relevance of changes in resource availability, measured here through intra and inter-specific competition, in explaining temporal trends in body mass, and demonstrate how they can operate across life-stages.

### Temporal trends in body mass

The consistent decrease in adult body mass of around 0.020 grams year^-1^ (0.11% year^-1^) in Wytham great tits adds to the growing body of evidence reporting declines in body mass and size across taxa (He et al., 2023; Teplitsky & Millien, 2014). This rate of change is similar in magnitude to other studies on birds in general (Yom-Tov et al., 2006; Youngflesh et al., 2022), for instance, body mass is declining at an average rate of 0.075% year^-1^ in 14 passerine species in England (Yom-Tov et al., 2006). This is also true when comparing the rate of change we report for Wytham to other great tit studies (Husby et al., 2011). As expected, and extending previous analyses (Garant et al., 2004), nestling body mass also decreased across our study period, at approximately 0.029 grams year^-1^ when considering all nestlings, and at 0.019 grams year^-1^ for the subset of nestlings that recruited to the breeding population. The rates of phenotypic change in adult and nestling mass in Wytham (−0.042 and −0.036 Haldanes, respectively) show a moderate to strong rate of change when compared to those reported across birds and other vertebrates (Gorné et al., 2025), highlighting the relevance of our results. For instance, the median absolute rate of change across bird traits is 0.004 Haldanes, and the median absolute rate of change for morphological traits across vertebrates is 0.010 Haldanes (see SM-8 and Gorné et al., 2025). A detailed analysis of the change in body mass, however, revealed a more complex temporal pattern, with a divergent direction of change between- and within-cohorts. While between cohorts the temporal trend was negative, within cohorts the change was positive, a pattern akin to Simpson’s Paradox. This suggests that the decline in mass found is the product of environmental changes at early life that carry over to adulthood. The pattern within cohorts likely reflects an increase in mass with age, a well described phenomenon in this and in other species (Furness & Robinson, 2019).

We found positive selection on nestling mass, slightly stronger to that previously reported by Garant et al. (2004) for nestling mass in Wytham between 1965 and 2000 (mean selection differential 0.209). This aligns with our temporal trends showing an increase in selection strength between 1978 and 2024 and suggests that the environmental conditions during the fledging and post-fledging phases may have become more challenging, potentially explaining the temporal trends in mass. Combined with the fact that, despite being heritable, the trends in adult mass are not aligned with a change at the genetic level, this strongly supports the conclusion that the observed trends represent a response to the change in the environmental conditions rather than evolution, an emerging general pattern when considering trends in body mass and size (Teplitsky & Millien, 2014).

### Temperature effects on body mass

Temporal trends in body mass have commonly been analysed in the light of climate change, with many studies showing declines in mass linked to warming, supporting the expectation from an intraspecific application of Bergmann’s rule (He et al., 2023; Teplitsky & Millien, 2014). However, the results from our two complementary approaches analysing the link between body mass and temperature do not support this mechanism in this case.

First, adult body mass was independent of the average temperature experienced during the breeding season (April to June) and the preceding winter (October to January), despite temperature in both periods having increased during the study period. By contrast, we found great tit nestlings to be heavier in warmer breeding seasons, something contrasting the trends generally found in adults. These positive effects of temperature are not unexpected and they align with those reported for hot extreme climatic events in our population (Satarkar, López-Idiáquez, et al., 2025). This positive link between temperature and body mass for nestlings might arise for several reasons. First, it might result from higher temperatures increasing resource availability by making prey more visible (McCarty & Winkler, 1999; Schöll et al., 2016). Moreover, warmer conditions can reduce direct and indirect thermoregulatory costs for nestlings, allowing them to allocate more energy to growth (Andreasson et al., 2018), while parents can spend more time foraging and less brooding (Dawson et al., 2005; Rodriguez & Barba, 2016).

Second, analysing the link between mass and temperature using specific time intervals for each adult individual and nestling brood showed a negative association in adults and a hump-shaped association in nestlings, with body mass being higher at intermediate temperatures. In adults, the decrease in mass at warmer temperatures can be associated with the effects of temperature on the predation-starvation trade-off (Cresswell et al., 2009; Houston & McNamara, 1993; Macleod et al., 2005). Under warmer environments, when foraging uncertainty is reduced, individuals lower their body mass to increase their flying efficiency which, in addition, also reduces the risks of being predated (Cresswell et al., 2009; Gosler et al., 1995). An alternative explanation is that the reduction in body mass at higher temperatures is a thermoregulatory response, as lighter individuals could, in theory, dissipate heat more effectively. Although this has long been the classic explanation for changes in body mass and size under warming (e.g. Youngflesh et al. 2022), recent work challenges this view because most reported changes in mass and size are unlikely to meaningfully impact heat dissipation (Nord et al., 2024). Thus, we argue that the former explanation, rather than the latter, best accounts for the negative association between temperature and adult great tit mass. In nestlings, the association between average temperature during development and mass was weaker than in adults and likely reflects temperature-driven variation in food availability. Temperature during breeding is tightly associated with the degree of synchrony with the phenology of winter moth caterpillars (López-Idiáquez et al., 2024), the main resource for great tit nestlings in our population (Hinks et al., 2015; Perrins, 1991). Broods raised at higher and lower temperatures experience a greater mismatch with the caterpillar peak, something that as shown here is associated with reduced nestling mass. This result is also consistent with our previous finding that breeding at high and low temperature imposes fitness costs to breeding adults (López-Idiáquez et al., 2024), further highlighting the importance of considering temperature at specific life-stages as a key modulator of breeding success.

### Resource availability effects on mass

An alternative explanation for the trends in body mass in both adult and nestlings is a temporal change in resource availability (Cox et al., 2019; Husby et al., 2011; Wilcox et al., 2024). We show that both resource availability proxies considered here, intra- and inter-specific competition and mismatch with the peak of food availability, impacted great tit adult and nestling mass.

For competition, our results show that great tit adult body mass showed weak but divergent associations with the number of great and blue tit pairs. While adult great tit body mass increased with a greater number of conspecifics at large spatial scales, it decreased with higher number of blue tits at small spatial scales. This divergence can be explained by the different competition forms of great tits and blue tits. While great tits are primarily interference competitors, blue tits are exploitation competitors (Dhondt, 2023; Minot, 1981). This means that while the effects of intraspecific competition manifests into smaller great tit territories at higher densities, the effects of interspecific competition with blue tits will arise through reduced resource availability in the territories of the great tits. These ideas, which align with our results, have received experimental support. For example studies have found that removing a subset of the great tit population increases the territory size of the remaining individuals (Both & Visser, 2000) and removing blue tits has a direct positive causal effect on the mass of the great tit nestlings breeding in the area (Minot, 1981). However, the strength of the associations we find between density at different spatial scales and great tit adult body mass are weak, probably reflecting population level processes linking resource availability and breeder density. For nestling mass, we found negative effects of both great and blue tit numbers, something aligning with previous results and reflecting an effect of both intra- and inter-specific competition (Both & Visser, 2000; Dhondt, 2023; Gamelon et al., 2019; Minot, 1981). Density effects of great tits were stronger across all spatial scales when compared to the effects of blue tits, where there was only one strong and significant negative effect on great tit nestling mass. These stronger and more spatially pervasive effects of great tit numbers on nestling mass are not surprising. Several studies in our and in other populations have shown that great tits have a competitive advantage over blue tits (Dhondt, 2011; Gamelon et al., 2019). Therefore, is likely that great tits are more sensitive to intra-than to inter-specific competition with blue tits as we have shown for body mass and has previously been shown for other processes such as population density regulation (Gamelon et al., 2019). This spatial variability in the results highlights that looking at the effects of competition on body mass at different spatial scales is important to develop a full understanding of how these two variables interact, a statement that can be extended to other environmental variables whose effects may vary spatially.

For mismatch with the peak of food availability, we found a weak but significant negative association with adult body mass, which is evident mostly due to a subset of individuals breeding early, and thus being mismatched with the peak of food availability. We suggest that this association is the product of the effects of mass on the timing of breeding, rather than the opposite, as this aligns with the well described pattern of heavier individuals laying earlier clutches in great tits and in other species (Descamps et al., 2011; Pigeault et al., 2020). For nestlings, maximum mass was achieved in those broods synchronized with the peak of food availability (i.e. mismatch ∼ 0) and then declined at both sides of the peak. This association can be explained by the highest availability of food for synchronous broods, which allows parents to more efficiently forage and feed their offspring (te Marvelde et al., 2011; Thomas et al., 2001). Overall, the reported association between nestling mass and mismatch adds to the large body of evidence highlighting the relevance of being synchronous with the peak of food availability for great tit and other species relying on ephemeral resources fitness (Simmonds et al., 2017; van Dis et al., 2023; Visser & Gienapp, 2019).

Taking the effects discussed above together, given that the number of great and blue tits breeding in Wytham has increased, and mismatch has remained stable across the study period, our results suggest that the temporal trend in body mass of nestlings is explained by an increase in intra- and inter-specific competition. This agrees with results of Husby et al. (2011) in highlighting the relevance of resource availability. However, in their case the trends in adult mass were explained by changes in mismatch and not great tit density. This divergence is likely caused by the different trends in the environmental variables, as for instance, while in Wytham great tits are successfully tracking the phenology of their prey and the opposite is true for the great tit populations studied in Husby et al. (2011). These two examples therefore illustrate that body mass can be shaped by different environmental drivers, even among the same species, reflecting population-specific responses to changes in ecological conditions.

### Carry over effects on adult body mass

Although adult body mass is sensitive to the variation in the three considered environmental factors, our results suggest that the marked decline in adult mass was not a direct response to them. This aligns with our analysis splitting the temporal trends in mass into the between- and within-cohort effects showing a negative temporal trend in the former but not the latter. Our results linking mass with several environmental factors suggest that the trends in adult mass are largely explained by developmental effects carrying-over to adulthood. Early-life effects are widely recognised as modulators of life-history trait expression during adulthood in bird and in other taxa (Brown et al., 2022; Monaghan, 2008; Pettorelli et al., 2002; Reid et al., 2003; Soma et al., 2006). For instance, they have been the main explanation for the decline in mass in a population of barn swallows *Hirundo rustica* in Italy (Romano et al., 2025). However, neither this nor most of the studies analysing temporal trends in mass tested the potential role of changes at early life as the explanation for the trends in adults, as we did. Although we lack a mechanistic explanation for the link between competition at early-life and adult mass, which could be explained by positive feedback loops or trade-offs, for instance (Fokkema et al., 2021; Kim et al., 2011), we provide evidence on the importance of considering environmental effects on different life-stages when explaining temporal trends in traits.

## Acknowledgements

We thank the numerous researchers and fieldworkers involved in the collection of the long-term data for the Wytham tit study over the past 79 years, and L. Cole for providing winter moth half-fall dates. We also thank S. R. Evans for his analytical and conceptual insight during the development of this work. The long-term population study has been supported by numerous funding sources, including recently by grants from BBSRC (BB/L006081/1), NERC (NE/K006274/1, NE/S010335/1) and a UKRI Frontiers Award (EP/X024520/1), which, together with NERC award NE/X000184/1 supported this work.

