## Supplementary material for "Not-so-great tit: early-life environment drives long-term decrease in adult body mass in a wild bird population": SM

**SM-1 Testing the effect of adult RFID identification on the temporal trends in body mass**

Since 2011, all captured great tits in Wytham Woods have been individually marked with an RFID tag, in addition to a metal ring, allowing individuals to be identified without physical recapture. Since then, we have used the RFID tags to identify breeding adults when they enter the nest boxes to feed their brood by placing an RFID reader at the nest entrance. The breeding adults identified using the RFIDs are not captured, which led to a decrease in the number of adults captured from 2011 onwards (SM-7, Fig. 1A), and thus their mass is not recorded. This reduction in the number of captured adults probably explains the larger standard errors shown in Fig. 1A of the main text between 2011 and 2024.

This approach will lead to an overrepresentation of dispersing individuals breeding in Wytham for the first time, in the dataset on adult body mass, as individuals lacking an RFID tag must be captured to be identified (and are then weighed). This could potentially lead to a bias in the reported temporal trends in adult mass, for instance, if this subset of individuals is different (i.e. higher or lower weights) than those born and/or already breeding in Wytham.

To analyse the potential biases introduced by the RFID adult identification we followed two complementary approaches. First, we analysed whether the body mass of individuals dispersing to Wytham and captured for the first time is different from that of individuals marked with an RFID tag, either as chicks if they were born in Wytham or during the first capture if they originated outside of the woods. We did this analysis both for the period 1978 – 2010, when all individuals were captured to be identified, and for the 2011 – 2024 period, when the sampling effort is biased towards individuals without the RFID tag. In both time periods, we found significant differences in body mass between the two groups, with individuals that had been born within, or previously captured as adults in Wytham being heavier than dispersers breeding in Wytham at the time of first capture as a breeding adult (period 1978 – 2010: 0.142±0.001 g, $F_{1,\ 13730.2}$=99.55, p<0.001; period 2011-2024: 0.105±0.032, $F_{1,\ 2225.1}$=11.26, <0.001, SM-7 Fig. 1B). Despite being significant, these differences are quite small, suggesting that the trends in adult body mass over the whole period reported in the main text are not a by-product of an overrepresentation of lighter individuals in recent years of the trend caused by the methodological shift.

We complemented this analysis by exploring the temporal trend before and after 2011, using the same model structure as in the main text. Our results show a significant decrease in adult body mass between 1978 and 2010 (-0.147±0.026, $F_{1,\ 33.8}$=31.12, p<0.001, SM-7, Fig. 1C). Moreover, the trend between 2011 and 2024 is also negative and of the same magnitude although not significant (-0.100±0.052, $F_{1,\ 12.16}$=3.656, p=0.079, SM-7, Fig. 1C). Note that in the models year is scaled to a mean of zero and standard deviation of unity. This, together with the gradual decrease in mass from year to year shown in Fig. 1A of the main text and in SM-7 Fig. 1C, confirms that the reported temporal trend is not a byproduct of the change in the way we identify breeding adults.

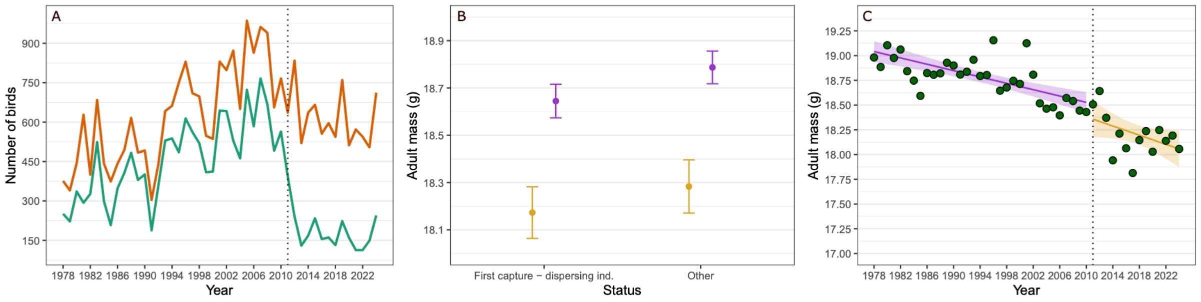


SM-7 Figure 1. A) Representation of the reduction in the number of great tit adults that are physically captured during the breeding seasons from 2011 onwards. The orange line is an estimate of the number of breeding individuals in the nest-boxes each year, calculated as the number of breeding pairs multiplied by 2. The green line represents the number of breeding adult individuals for which body mass data is available each year. B) Predicted mean adult breeding mass ± 95% credible interval differences between dispersing individuals captured for the first time and the rest of the population. In purple the period from 1978 to 2010 and in yellow the period from 2011 to 2024. C) Temporal trends in adult body mass; note the difference in scale in parts B and C. The purple line and ribbon represent the trend predicted between 1978 and 2010, and yellow line and ribbon the predicted trend between 2011 and 2024. Green dots represent the mean adult mass each year.

### SM-2: Results from the random slope model

SM8-Table 1: Results from the random slope model testing the temporal trends in body mass within and between cohorts. Year in the random slope and as a fixed effect represents a continuous temporal trend. Year and year of birth as random effects represent a group-level effect (i.e. categorical variables). The subscripts in section represent the section names in Wytham woods. Bold denotes estimates whose 95% credible intervals (CI) did not include 0.

| | Estimate | 95% CI |
|---|---|---|
| Adult body mass (n=17785) | | |
| *Random effects* | | |
| **Intercept** | **1.338** | **[0.874, 1.799]** |
| **Year*Year of birth** | **0.385** | **[0.041, 0.690]** |
| $Cor_{Int\text{-}year}$ | 0.015 | [-0.531, 0.510] |
| **Ring** | **0.651** | **[0.636, 0.665]** |
| **Year** | **0.140** | **[0.105, 0.185]** |
| *Fixed effects* | | |
| **Intercept** | **18.412** | **[17.959, 18.822]** |
| **Year** | **0.833** | **[0.524, 1.090]** |
| **$Sex_{male}$** | **0.361** | **[0.331, 0.389]** |
| $Section_{c}$ | -0.063 | [-0.128, 0.002] |
| **$Section_{cp}$** | **0.098** | **[0.028, 0.161]** |
| **$Section_{ex}$** | **0.056** | **[0.026, 0.004]** |
| $Section_{mp}$ | -0.032 | [-0.096, 0.029] |
| $Section_{o}$ | -0.035 | [-0.090, 0.022] |
| **$Section_{p}$** | **-0.153** | **[-0.257, -0.054]** |
| $Section_{sw}$ | -0.051 | [-0.110, 0.005] |
| **$Section_{w}$** | **0.134** | **[0.078, 0.191]** |

**SM-3: Temporal trends in selection differentials on nestling body mass**

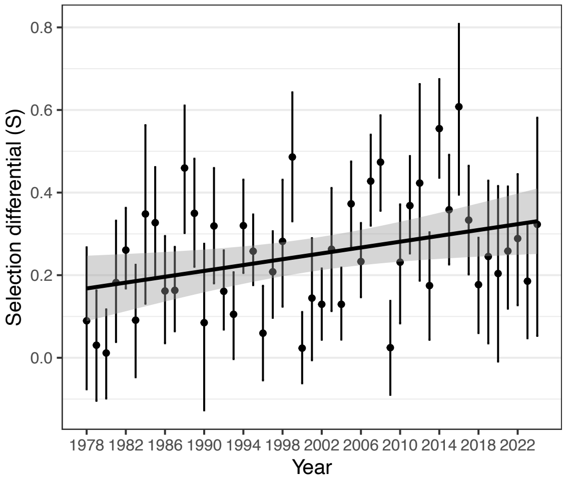


SM-6 Fig. 1: Temporal trends in yearly selection differentials on nestling mass. Dots represent the selection differential and whiskers the 95% confidence intervals (obtained from 1000 bootstraps).

**SM-4: Temporal trends in the breeding values**

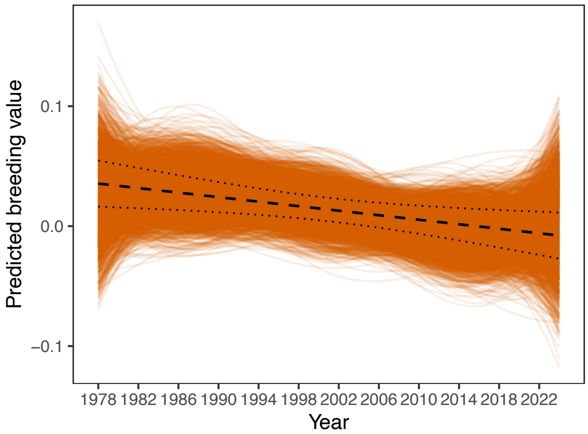


SM-2 Figure 2: Temporal trends in adult mass breeding values. Black dashed line represents the linear regression of posterior mean breeding value against mean breeding year and the 95% confidence interval (dotted lines). Orange lines represent the change in the breeding values allowing for a non-linear change using splines obtained from each posterior sample from the Bayesian posterior distribution.

**SM-5: Association between temperature and body mass**

SM-5 Table 1: Results from the linear mixed model linking temperature in spring (April to June) and adult body mass. Bold denotes statistical significance.

| | **Estimate** | **SE** | ***F*** | ***P*** |
|---|---|---|---|---|
| Adult body mass (n=17785) | | | | |
| Temperature spring | -0.015 | 0.027 | $F_{1,45.5}$=0.317 | 0.575 |
| **Section** | | | **$F_{8, 13309.9}$=10.86** | **<0.001** |
| **Year** | **-0.222** | **0.025** | **$F_{1,48.6}$=74.96** | **<0.001** |
| **$Sex_{male}$** | **0.358** | **0.015** | **$F_{1,11604.2}$=557.5** | **<0.001** |
| **$Age_{juv}$** | **-0.256** | **0.011** | **$F_{1,12556.8}$=520.2** | **<0.001** |

SM-5 Table 2: Results from the linear mixed model linking temperature in winter (October to January) and adult body mass. Bold denotes statistical significance.

| | **Estimate** | **SE** | ***F*** | ***P*** |
|---|---|---|---|---|
| Adult body mass (n=17785) | | | | |
| Temperature winter | -0.011 | 0.027 | F1, 43.2=0.001 | 0.976 |
| **Section** | | | **$F_{8, 13310.1}$=10.87** | **<0.001** |
| **Year** | **-0.230** | **0.024** | **$F_{1, 47.4}$=88.59** | **<0.001** |
| **$Sex_{male}$** | **0.358** | **0.015** | **$F_{1,11604.5}$=557.5** | **<0.001** |
| **$Age_{juv}$** | **-0.256** | **0.011** | **$F_{1,12555.1}$=520.5** | **<0.001** |

SM-5 Table 3: Results from the linear mixed model linking temperature in spring (April to June) and nestling body mass. Bold denotes statistical significance.

| | **Estimate** | **SE** | ***F*** | ***P*** |
|---|---|---|---|---|
| Nestling body mass (n=77787) | | | | |
| **Temperature winter** | **0.185** | **0.071** | **$F_{1, 44.10}$=6.727** | **0.012** |
| **Section** | | | **$F_{8, 829.30}$=27.89** | **<0.001** |
| **Year** | **-0.474** | **0.071** | **$F_{1, 44.10}$=44.28** | **<0.001** |

### SM-6: Association between breeding density and body mass

SM-6 Table 1: Results from the linear mixed models linking breeding adult body mass and the number of great tit pairs breeding at different distance buffers.

| | Estimate | SE | *F* | *P* |
|---|---|---|---|---|
| | Adult body mass (n=17785) | | | |
| | 100 m. buffer | | | |
| Nº great tits | 0.005 | 0.007 | $F_{1,15634.1}$=0.518 | 0.471 |
| **Section** | | | **$F_{8,13568.6}$=10.826** | **<0.001** |
| **Year** | **-0.231** | **0.020** | **$F_{1,50.6}$=127.12** | **<0.001** |
| **$Sex_{male}$** | **0.358** | **0.151** | **$F_{1, 11603.9}$=557.68** | **<0.001** |
| **$Age_{juv}$** | **-0.256** | **0.011** | **$F_{1, 12552.9}$=520.71** | **<0.001** |
| | 200 m. buffer | | | |
| Nº great tits | -0.003 | 0.009 | $F_{1, 13508.9}$=0.114 | 0.735 |
| **Section** | | | **$F_{8,13627.6}$=10.87** | **<0.001** |
| **Year** | **-0.231** | **0.020** | **$F_{1,50.4}$=123.80** | **<0.001** |
| **$Sex_{male}$** | **0.358** | **0.015** | **$F_{1, 11604.5}$=557.41** | **<0.001** |
| **$Age_{juv}$** | **-0.256** | **0.011** | **$F_{1, 12551.1}$=520.36** | **<0.001** |
| | 300 m. buffer | | | |
| Nº great tits | 0.001 | 0.009 | $F_{1, 10696.6}$=0.016 | 0.897 |
| **Section** | | | **$F_{8, 13569.9}$=10.837** | **<0.001** |
| **Year** | **-0.231** | **0.020** | **$F_{1, 50.4}$=125.40** | **<0.001** |
| **$Sex_{male}$** | **0.358** | **0.015** | **$F_{1,11604.3}$=557.54** | **<0.001** |
| **$Age_{juv}$** | **-0.256** | **0.011** | **$F_{1, 12554.5}$=520.52** | **<0.001** |
| | 400 m. buffer | | | |
| Nº great tits | -0.001 | 0.009 | $F_{1, 8639.5}$=0.031 | 0.859 |
| **Section** | | | **$F_{8, 13496.2}$=10.76** | **<0.001** |
| **Year** | **-0.231** | **0.020** | **$F_{1, 50.3}$=123.82** | **<0.001** |
| **$Sex_{male}$** | **0.358** | **0.015** | **$F_{1,11603.9}$=557.49** | **<0.001** |
| **$Age_{juv}$** | **-0.256** | **0.011** | **$F_{1, 12554.5}$=520.586** | **<0.001** |
| | 500 m. buffer | | | |
| Nº great tits | 0.003 | 0.010 | $F_{1,6948.8}$=0.116 | 0.733 |
| **Section** | | | **$F_{8, 13467.5}$=10.34** | **<0.001** |
| **Year** | **-0.231** | **0.020** | **$F_{1, 50.3}$=126.317** | **<0.001** |
| **$Sex_{male}$** | **0.358** | **0.015** | **$F_{1, 11602.9}$=557.56** | **<0.001** |
| **$Age_{juv}$** | **-0.256** | **0.011** | **$F_{1, 12557.8}$=520.20** | **<0.001** |
| | 600 m. buffer | | | |
| Nº great tits | 0.007 | 0.010 | $F_{1, 5920.0}$=0.547 | 0.459 |
| **Section** | | | **$F_{8, 13499.6}$=0.855** | **<0.001** |
| **Year** | **-0.232** | **0.020** | **$F_{1, 50.5}$=128.49** | **<0.001** |
| **$Sex_{male}$** | **0.358** | **0.151** | **$F_{1, 11603.1}$=557.48** | **<0.001** |
| **$Age_{juv}$** | **-0.256** | **0.011** | **$F_{1, 12557.1}$=519.9** | **<0.001** |
| | 700 m. buffer | | | |
| Nº great tits | 0.0005 | 0.011 | $F_{1, 4892.6}$=0.002 | 0.959 |
| **Section** | | | **$F_{8, 13464.6}$=9.923** | **<0.001** |
| **Year** | **-0.231** | **0.020** | **$F_{1, 50.2}$=124.55** | **<0.001** |
| **$Sex_{male}$** | **0.358** | **0.015** | **$F_{1,11603.0}$=557.51** | **<0.001** |
| **$Age_{juv}$** | **-0.256** | **0.011** | **$F_{1,12554.2}$=520.48** | **<0.001** |
| | 800 m. buffer | | | |
| Nº great tits | -0.006 | 0.012 | $F_{1, 3822.0}$=0.285 | 0.593 |
| **Section** | | | **$F_{8,13437.8}$=10.14** | **<0.001** |
| **Year** | **-0.230** | **0.020** | **$F_{1,49.9}$=120.47** | **<0.001** |
| **$Sex_{male}$** | **0.358** | **0.015** | **$F_{1, 11602.3}$=557.56** | **<0.001** |
| **$Age_{juv}$** | **-0.256** | **0.011** | **$F_{1, 12551.1}$=520.79** | **<0.001** |
| | 900 m. buffer | | | |
| Nº great tits | **-0.009** | **-0.013** | **$F_{1, 2806.2}$=0.539** | **0.462** |
| **Section** | | | **$F_{8, 12908.2}$=10.268** | **<0.001** |
| **Year** | **0.229** | **0.021** | **$F_{1, 49.7}$=118.190** | **<0.001** |
| **$Sex_{male}$** | **0.358** | **0.015** | **$F_{1,11602.0}$=557.59** | **<0.001** |
| **$Age_{juv}$** | **-0.256** | **0.011** | **$F_{1, 12547.9}$=520.79** | **<0.001** |

| | | | | |
|---|---|---|---|---|
| 1000 m. buffer | | | | |
| Nº great tits | -0.010 | 0.014 | $F_{1,2078.8}$=0.526 | 0.468 |
| **Section** | | | **$F_{8, 12408}$=10.358** | **<0.001** |
| **Year** | **-0.229** | **0.021** | **$F_{1, 49.7}$=117.22** | **<0.001** |
| **Sex$_{male}$** | **0.358** | **0.015** | **$F_{1, 11601.7}$=557.57** | **<0.001** |
| **Age$_{juv}$** | **-0.256** | **0.011** | **$F_{1, 12545.5}$=520.68** | **<0.001** |
| 1250 m. buffer | | | | |
| Nº great tits | **-0.012** | **0.015** | **$F_{1,1573.6}$=0.644** | **0.422** |
| **Section** | | | **$F_{8, 11931.3}$=10.81** | **<0.001** |
| **Year** | **-0.022** | **0.021** | **$F_{1, 49.8}$=115.14** | **<0.001** |
| **Sex$_{male}$** | **0.358** | **0.015** | **$F_{1,11602.7}$=557.60** | **<0.001** |
| **Age$_{juv}$** | **-0.256** | **0.011** | **$F_{1, 12547.1}$=520.77** | **<0.001** |
| 1500 m. buffer | | | | |
| Nº great tits | -0.009 | 0.015 | $F_{1, 1772.1}$=0375 | 0.540 |
| **Section** | | | **$F_{8,12216.3}$=10.91** | **<0.001** |
| **Year** | **-0.229** | **0.021** | **$F_{1,50.1}$=11.08** | **<0.001** |
| **Sex$_{male}$** | **-0.358** | **0.015** | **$F_{1, 11603.1}$=557.56** | **<0.001** |
| **Age$_{juv}$** | **-0.256** | **0.011** | **$F_{1,12549.5}$=520.71** | **<0.001** |
| 1750 m. buffer | | | | |
| Nº great tits | 0.011 | 0.015 | $F_{1, 1389.7}$=0.542 | 0.461 |
| **Section** | | | **$F_{8, 11651.6}$=10.93** | **<0.001** |
| **Year** | **-0.233** | **0.020** | **$F_{1,51.3}$=128.16** | **<0.001** |
| **Sex$_{male}$** | **0.358** | **0.015** | **$F_{1,11603.6}$=557.47** | **<0.001** |
| **Age$_{juv}$** | **-0.256** | **0.020** | **$F_{1, 12554.4}$=520.50** | **<0.001** |
| 2000 m. buffer | | | | |
| Nº great tits | -0.029 | 0.016 | F1, 744.2=3.00 | 0.083 |
| **Section** | | | **$F_{8, 10151.3}$=11.231** | **<0.001** |
| **Year** | **-0.237** | **0.020** | **$F_{1, 52.4}$=137.36** | **<0.001** |
| **Sex$_{male}$** | **0.357** | **0.015** | **$F_{1, 11603.9}$=557.37** | **<0.001** |
| **Age$_{juv}$** | **-0.256** | **0.011** | **$F_{1, 12556.1}$=520.75** | **<0.001** |
| 2500 m. buffer | | | | |
| Nº great tits | 0.036 | 0.019 | $F_{1, 198.6}$=3.261 | 0.072 |
| **Section** | | | **$F_{8, 6218.1}$=11.27** | **<0.001** |
| **Year** | **-0.241** | **0.020** | **$F_{1, 53.3}$=139.00** | **<0.001** |
| **Sex$_{male}$** | **0.357** | **0.015** | **$F_{1, 11604.8}$=557.15** | **<0.001** |
| **Age$_{juv}$** | **-0.256** | **0.011** | **$F_{1, 12554.5}$=520.95** | **<0.001** |
| 3000 m. buffer | | | | |
| **Nº great tits** | **0.048** | **0.021** | **$F_{1,95.1}$=4.91** | **0.028** |
| **Section** | | | **$F_{8, 8390.0}$=11.18** | **<0.001** |
| **Year** | **-0.246** | **0.020** | **$F_{1, 54.0}$=143.10** | **<0.001** |
| **Sex$_{male}$** | **0.357** | **0.015** | **$F_{1, 11605.6}$=556.98** | **<0.001** |
| **Age$_{juv}$** | **-0.256** | **0.011** | **$F_{1, 12555.5}$=520.78** | **<0.001** |
| 4000 m. buffer | | | | |
| **Nº great tits** | **0.072** | **0.025** | **$F_{1, 44.6}$=8.010** | **0.006** |
| **Section** | | | **$F_{8, 13311.5}$=10.86** | **<0.001** |
| **Year** | **-0.254** | **0.021** | **F1, 51.4=146.96** | **<0.001** |
| **Sex$_{male}$** | **0.357** | **0.015** | **$F_{1, 11605.7}$=557.13** | **<0.001** |
| **Age$_{juv}$** | **-0.256** | **0.011** | **$F_{1, 12555.5}$=520.17** | **<0.001** |

SM-6 Table 2: Results from the linear mixed models linking breeding adult body mass and the number of blue tit pairs breeding at different distance buffers.

| | Estimate | SE | *F* | *P* |
|---|---|---|---|---|
| | | Adult body mass (n=17785) | | |
| | | 100 m. buffer | | |
| Nº blue tits | -0.019 | 0.007 | $F_{1, 16837.0}$=6.581 | 0.010 |
| **Section** | | | **$F_{8, 13603.5}$=10.93** | **<0.001** |
| **Year** | **-0.229** | **0.020** | **$F_{1,50.7}$=122.70** | **<0.001** |
| **$Sex_{male}$** | **0.357** | **0.015** | **$F_{1,11604.2}$=557.27** | **<0.001** |
| **$Age_{juv}$** | **-0.256** | **0.011** | **$F_{1,12556.0}$=521.32** | **<0.001** |
| | | 200 m. buffer | | |
| Nº blue tits | **-0.022** | **-0.008** | **$F_{1, 17339.2}$=7.253** | **0.007** |
| **Section** | | | **$F_{8, 13632.6}$=10.95** | **<0.001** |
| **Year** | **-0.228** | **0.020** | **$F_{1,50.8}$=121.22** | **<0.001** |
| **$Sex_{male}$** | **0.357** | **0.015** | **$F_{1,11602.8}$=557.14** | **<0.001** |
| **$Age_{juv}$** | **0.256** | **0.011** | **$F_{1,12546.2}$=520.22** | **<0.001** |
| | | 300 m. buffer | | |
| Nº blue tits | **-0.021** | **0.008** | **$F_{1,16994.6}$=6.095** | **0.013** |
| **Section** | | | **$F_{8, 13644.5}$=10.95** | **<0.001** |
| **Year** | **-0.227** | **0.020** | **$F_{1, 51.1}$=120.17** | **<0.001** |
| **$Sex_{male}$** | **0.358** | **0.015** | **$F_{1, 11601.3}$=557.43** | **<0.001** |
| **$Age_{juv}$** | **-0.256** | **0.011** | **$F_{1, 12547.0}$=520.83** | **<0.001** |
| | | 400 m. buffer | | |
| Nº blue tits | **-0.019** | **0.008** | **$F_{1, 16417.6}$=5.117** | **0.023** |
| **Section** | | | **$F_{8, 13610.7}$=11.107** | **<0.001** |
| **Year** | **-0.227** | **0.020** | **$F_{1,51.4}$=119.233** | **<0.001** |
| **$Sex_{male}$** | **0.358** | **0.015** | **$F_{1, 11601.0}$=557.45** | **<0.001** |
| **$Age_{juv}$** | **-0.256** | **0.011** | **$F_{1,11601.0}$=520.12** | **<0.001** |
| | | 500 m. buffer | | |
| Nº blue tits | **-0.017** | **0.008** | **$F_{1, 15663.3}$=3.867** | **0.049** |
| **Section** | | | **$F_{8, 13558.7}$=11.19** | **<0.001** |
| **Year** | **-0.227** | **0.020** | **$F_{1,51.7}$=119.07** | **<0.001** |
| **$Sex_{male}$** | **0.358** | **0.015** | **$F_{1,11602.2}$=557.44** | **<0.001** |
| **$Age_{juv}$** | **-0.256** | **0.011** | **$F_{1,12549.5}$=520.56** | **<0.001** |
| | | 600 m. buffer | | |
| Nº blue tits | -0.016 | 0.008 | $F_{1, 14818.0}$=3.413 | 0.064 |
| **Section** | | | **$F_{8, 13521.0}$=11.25** | **<0.001** |
| **Year** | **-0.226** | **0.020** | **$F_{1, 52.0}$=118.51** | **<0.001** |
| **$Sex_{male}$** | **0.358** | **0.015** | **$F_{1, 11603.0}$=557.73** | **<0.001** |
| **$Age_{juv}$** | **-0.256** | **0.011** | **$F_{1, 12552.0}$=520.22** | **<0.001** |
| | | 700 m. buffer | | |
| Nº bue tits | **-0.019** | **0.008** | **$F_{1, 13569.6}$=4.696** | **0.030** |
| **Section** | | | **$F_{8, 13528.0}$=11.44** | **<0.001** |
| **Year** | **-0.225** | **0.020** | **$F_{1, 52.3}$=116.32** | **<0.001** |
| **$Sex_{male}$** | **0.358** | **0.015** | **$F_{1, 11602.1}$=557.93** | **<0.001** |
| **$Age_{juv}$** | **-0.256** | **0.011** | **$F_{1, 12553.0}$=519.80** | **<0.001** |
| | | 800 m. buffer | | |
| Nº blue tits | **-0.024** | **0.009** | **$F_{1, 11772.8}$=6.984** | **0.008** |
| **Section** | | | **$F_{8, 13541.0}$=11.71** | **<0.001** |
| **Year** | **-0.222** | **0.020** | **$F_{1, 52.8}$=11.87** | **<0.001** |
| **$Sex_{male}$** | **0.358** | **0.015** | **$F_{1, 11601.6}$=558.08** | **<0.001** |
| **$Age_{juv}$** | **-0.256** | **0.011** | **$F_{1, 12554.4}$=519.47** | **<0.001** |
| | | 900 m. buffer | | |
| Nº blue tits | **-0.027** | **0.010** | **$F_{1, 9741.1}$=7.286** | **0.006** |
| **Section** | | | **$F_{8, 13547.8}$=11.77** | **<0.001** |
| **Year** | **-0.221** | **0.021** | **$F_{1, 53.5}$=110.34** | **<0.001** |
| **$Sex_{male}$** | **0.358** | **0.015** | **$F_{1, 11601.3}$=558.26** | **<0.001** |
| **$Age_{juv}$** | **-0.256** | **0.011** | **$F_{1, 12553.9}$=518.66** | **<0.001** |
| | | 1000 m. buffer | | |
| Nº blue tits | **-0.026** | **0.010** | **$F_{1, 7918.4}$=6.135** | **0.013** |

| | | | | |
|---|---|---|---|---|
| **Section** | | | **$F_{8. 13544.9}$=11.63** | **<0.001** |
| **Year** | **-0.221** | **0.021** | **$F_{1, 54.3}$=109.07** | **<0.001** |
| **$Sex_{male}$** | **0.358** | **0.015** | **$F_{1, 11601.2}$=558.24** | **<0.001** |
| **$Age_{juv}$** | **-0.256** | **0.011** | **$F_{1, 12.555.0}$=518.38** | **<0.001** |
| 1250 m. buffer | | | | |
| Nº blue tits | -0.020 | 0.012 | $F_{1, 5445.5}$=2.842 | 0.091 |
| **Section** | | | **$F_{8, 13398.1}$=11.22** | **<0.001** |
| **Year** | **-0.222** | **0.021** | **$F_{1, 56.0}$=109.30** | **<0.001** |
| **$Sex_{male}$** | **0.358** | **0.015** | **$F_{1, 11600.8}$=558.17** | **<0.001** |
| **$Age_{juv}$** | **-0.256** | **0.011** | **$F_{1, 12555.2}$=519.42** | **<0.001** |
| 1500 m. buffer | | | | |
| Nº blue tits | -0.015 | 0.012 | $F_{1, 5493.4}$=1.475 | 0.224 |
| **Section** | | | **$F_{8, 13359.6}$=11.00** | **<0.001** |
| **Year** | **-0.225** | **0.021** | **$F_{1, 56.2}$=111.57** | **<0.001** |
| **$Sex_{male}$** | **0.358** | **0.015** | **$F_{1, 11601.8}$=557.78** | **<0.001** |
| **$Age_{juv}$** | **-0.256** | **0.011** | **$F_{1, 12554.9}$=519.89** | **<0.001** |
| 1750 m. buffer | | | | |
| Nº blue tits | 0.001 | 0.013 | $F_{1, 4835.1}$=0.016 | 0.897 |
| **Section** | | | **$F_{8, 13206.8}$=10.84** | **<0.001** |
| **Year** | **-0.230** | **0.021** | **$F_{1, 56.7}$=116.87** | **<0.001** |
| **$Sex_{male}$** | **0.358** | **0.014** | **$F_{1, 11602.2}$=557.53** | **<0.001** |
| **$Age_{juv}$** | **-0.256** | **0.011** | **$F_{1, 12551.3}$=520.36** | **<0.001** |
| 2000 m. buffer | | | | |
| Nº blue tits | -0.010 | 0.014 | $F_{1,2779.9}$=0.472 | 0.492 |
| **Section** | | | **$F_{8, 12670.7}$=10.92** | **<0.001** |
| **Year** | **-0.235** | **0.021** | **$F_{1, 58.9}$=118.91** | **<0.001** |
| **$Sex_{male}$** | **0.358** | **0.015** | **$F_{1, 11603.0}$=557.49** | **<0.001** |
| **$Age_{juv}$** | **-0.256** | **0.011** | **$F_{1, 12550.3}$=520.97** | **<0.001** |
| 2500 m. buffer | | | | |
| **Nº blue tits** | **0.037** | **0.018** | **$F_{1, 571.5}$=4.07** | **0.043** |
| **Section** | | | **$F_{8, 9086.5}$=11.346** | **<0.001** |
| **Year** | **-0.250** | **0.022** | **$F_{1, 64.7}$=119.09** | **<0.001** |
| **$Sex_{male}$** | **0.358** | **0.015** | **$F_{1, 11605.6}$=557.79** | **<0.001** |
| **$Age_{juv}$** | **-0.257** | **0.011** | **$F_{1, 12545.9}$=522.42** | **<0.001** |
| 3000 m. buffer | | | | |
| Nº blue tits | -0.019 | 0.020 | $F_{1,200.5}$=0.868 | 0.352 |
| **Section** | | | **$F_{8, 7302.4}$=10.89** | **<0.001** |
| **Year** | **-0.243** | **-0.024** | **$F_{1,66.3}$=100.34** | **<0.001** |
| **$Sex_{male}$** | **0.358** | **0.015** | **$F_{1,11605.0}$=557.63** | **<0.001** |
| **$Age_{juv}$** | **-0.256** | **0.011** | **$F_{1,12551.2}$=521.12** | **<0.001** |
| 4000 m. buffer | | | | |
| Nº blue tits | 0.004 | 0.028 | $F_{1, 46.2}$=0.030 | 0.863 |
| **Section** | | | **$F_{8, 13309.8}$=10.87** | **<0.001** |
| **Year** | **-0.234** | **0.027** | **$F_{1,47.7}$=71.54** | **<0.001** |
| **$Sex_{male}$** | **0.358** | **0.015** | **$F_{1,11604.4}$=557.5** | **<0.001** |
| **$Age_{juv}$** | **-0.256** | **0.011** | **$F_{1,12553.0}$=520.61** | **<0.001** |

SM-6 Table 3: Results from the linear mixed models linking breeding nestling body mass and the number of great tit pairs breeding at different distance buffers.

| | Estimate | SE | *F* | *P* |
|---|---|---|---|---|
| Adult body mass (n=77787) | | | | |
| 100 m. buffer | | | | |
| Nº great tits | **-0.092** | **0.016** | $\mathbf{F_{1,5352.1}=30.41}$ | **<0.001** |
| **Section** | | | $\mathbf{F_{8,841.4}=31.85}$ | **<0.001** |
| **Year** | **-0.372** | **0.064** | $\mathbf{F_{1,45.5}=32.98}$ | **<0.001** |
| 200 m. buffer | | | | |
| **Nº great tits** | **-0.044** | **0.019** | $\mathbf{F_{1,3145.75}=5.360}$ | **0.020** |
| **Section** | | | $\mathbf{F_{8,839.32}=31.140}$ | **<0.001** |
| **Year** | **-0.376** | **0.064** | $\mathbf{F_{1,45.58}=33.82}$ | **<0.001** |
| 300 m. buffer | | | | |
| Nº great tits | -0.005 | 0.020 | $F_{1,2439.12}=0.062$ | 0.802 |
| **Section** | | | $\mathbf{F_{8,815.17}=31.28}$ | **<0.001** |
| **Year** | **-0.381** | **0.064** | $\mathbf{F_{1,45.78}=34.94}$ | **<0.001** |
| 400 m. buffer | | | | |
| Nº great tits | 0.009 | 0.020 | $F_{1,1914.03}=0.203$ | 0.652 |
| **Section** | | | $\mathbf{F_{8,796.55}=31.97}$ | **<0.001** |
| **Year** | **-0.382** | **0.064** | $\mathbf{F_{1,45.68}=35.27}$ | **<0.001** |
| 500 m. buffer | | | | |
| Nº great tits | -0.0007 | 0.021 | $F_{1,1728.67}=0.001$ | 0.971 |
| **Section** | | | $\mathbf{F_{8,796.95}=31.90}$ | **<0.001** |
| **Year** | **-0.382** | **0.064** | $\mathbf{F_{1,45.70}=34.92}$ | **<0.001** |
| 600 m. buffer | | | | |
| Nº great tits | -0.016 | 0.022 | $F_{1,1559.6}=0.543$ | 0.461 |
| **Section** | | | $\mathbf{F_{8,807.18}=31.66}$ | **<0.001** |
| **Year** | **-0.379** | **0.064** | $\mathbf{F_{1,45.73}=34.36}$ | **<0.001** |
| 700 m. buffer | | | | |
| Nº great tits | -0.034 | 0.023 | $F_{1,1586.28}=2.129$ | 0.144 |
| **Section** | | | $\mathbf{F_{8,827.81}=31.43}$ | **<0.001** |
| **Year** | **-0.375** | **0.064** | $\mathbf{F_{1,46.08}=33.80}$ | **<0.001** |
| 800 m. buffer | | | | |
| **Nº great tits** | **-0.054** | **0.025** | $\mathbf{F_{1,1710.98}=4.428}$ | **0.035** |
| **Section** | | | $\mathbf{F_{8,848.13}=31.30}$ | **<0.001** |
| **Year** | **-0.371** | **0.064** | $\mathbf{F_{1,45.91}=32.82}$ | **<0.001** |
| 900 m. buffer | | | | |
| **Nº great tits** | **-0.071** | **0.028** | $\mathbf{F_{1,1863.74}=6.37}$ | **0.011** |
| **Section** | | | $\mathbf{F_{8,867.88}=31.35}$ | **<0.001** |
| **Year** | **-0.367** | **0.065** | $\mathbf{F_{1,46.03}=31.35}$ | **<0.001** |
| 1000 m. buffer | | | | |
| **Nº great tits** | **-0.090** | **0.030** | $\mathbf{F_{1,1971.39}=8.88}$ | **0.002** |
| **Section** | | | $\mathbf{F_{8,874.63}=31.82}$ | **<0.001** |
| **Year** | **-0.362** | **0.065** | $\mathbf{F_{1,46.15}=30.85}$ | **<0.011** |
| 1250 m. buffer | | | | |
| **Nº great tits** | **-0.175** | **0.033** | $\mathbf{F_{1,1821.70}=28.18}$ | **<0.001** |
| **Section** | | | $\mathbf{F_{8,871.52}=36.02}$ | **<0.001** |
| **Year** | **-0.341** | **0.066** | $\mathbf{F_{1,46.04}=26.19}$ | **<0.001** |
| 1500 m. buffer | | | | |
| **Nº great tits** | **-0.185** | **0.032** | $\mathbf{F_{1,1566.95}=33.17}$ | **<0.001** |
| **Section** | | | $\mathbf{F_{8,848.29}=37.026}$ | **<0.001** |
| **Year** | **-0.339** | **0.066** | $\mathbf{F_{1,45.98}=25.74}$ | **<0.001** |
| 1750 m. buffer | | | | |
| **Nº great tits** | **-0.172** | **0.032** | $\mathbf{F_{1,1362.22}=27.387}$ | **<0.001** |
| **Section** | | | $\mathbf{F_{8,844.46}=35.98}$ | **<0.001** |
| **Year** | **-0.341** | **0.066** | $\mathbf{F_{1,46.12}=26.21}$ | **<0.001** |
| 2000 m. buffer | | | | |
| **Nº great tits** | **-0.158** | **0.036** | $\mathbf{F_{1,1189.28}=18.69}$ | **<0.001** |
| **Section** | | | $\mathbf{F_{8,861.43}=34.35}$ | **<0.001** |
| **Year** | **-0.342** | **0.066** | $\mathbf{F_{1,36.34}=26.38}$ | **<0.001** |
| 2500 m. buffer | | | | |
| **Nº great tits** | **-0.122** | **0.045** | $\mathbf{F_{1,470.57}=7.103}$ | **0.007** |

| | | | | |
|---|---|---|---|---|
| **Section** | | | $\mathbf{F_{8,861.21}=31.71}$ | **<0.001** |
| **Year** | **-0.345** | **0.067** | $\mathbf{F_{1,47.20}=26.36}$ | **<0.001** |
| | | 3000 m. buffer | | |
| **Nº great tits** | **-0.161** | **0.055** | $\mathbf{F_{1,180.44}=8.536}$ | **0.003** |
| **Section** | | | $\mathbf{F_{8,841.81}=32.01}$ | **<0.001** |
| **Year** | **-0.327** | **0.070** | $\mathbf{F_{1,45.18}=21.88}$ | **<0.001** |
| | | 4000 m. buffer | | |
| Nº great tits | -0.010 | 0.074 | $F_{1,43.93}=0.019$ | 0.889 |
| **Section** | | | $\mathbf{F_{8,817.53}=31.93}$ | **<0.001** |
| **Year** | **-0.378** | **0.070** | $\mathbf{F_{1,44.21}=28.71}$ | **<0.00§** |

SM-6 Table 4: Results from the linear mixed models linking breeding nestling body mass and the number of blue tit pairs breeding at different distance buffers.

| | Estimate | SE | *F* | *P* |
|---|---|---|---|---|
| | | Adult body mass (n=77787) | | |
| | | 100 m. buffer | | |
| Nº blue tits | -0.027 | 0.016 | $F_{1,5461.2}=2.753$ | 0.097 |
| **Section** | | | $\mathbf{F_{8,853.5}=31.325}$ | **<0.001** |
| **Year** | **-0.378** | **0.064** | $\mathbf{F_{1,45.5}=34.44}$ | **<0.001** |
| | | 200 m. buffer | | |
| Nº blue tits | -0.018 | 0.017 | $F_{1,4207.9}=1.083$ | 0.298 |
| **Section** | | | $\mathbf{F_{8,850.5}=31.309}$ | **<0.001** |
| **Year** | **-0.379** | **0.064** | $\mathbf{F_{1,45.6}=34.48}$ | **<0.001** |
| | | 300 m. buffer | | |
| Nº blue tits | -0.004 | 0.018 | $F_{1,3830.0}=0.065$ | 0.798 |
| **Section** | | | $\mathbf{F_{8,841.6}=31.38}$ | **<0.001** |
| **Year** | **-0.381** | **0.064** | $\mathbf{F_{1,45.8}=34.80}$ | **<0.001** |
| | | 400 m. buffer | | |
| Nº blue tits | 0.015 | 0.018 | $F_{1,3404.0}=0.727$ | 0.393 |
| **Section** | | | $\mathbf{F_{8,830.7}=32.03}$ | **<0.001** |
| **Year** | **-0.386** | **0.064** | $\mathbf{F_{1,46.0}=35.61}$ | **<0.001** |
| | | 500 m. buffer | | |
| Nº blue tits | 0.030 | 0.018 | $F_{1,2983.26}=2.685$ | 0.101 |
| **Section** | | | $\mathbf{F_{8,824.89}=32.48}$ | **<0.001** |
| **Year** | **-0.392** | **0.065** | $\mathbf{F_{1,46.11}=36.38}$ | **<0.001** |
| | | 600 m. buffer | | |
| Nº blue tits | 0.037 | 0.018 | $F_{1,2612.43}=3.879$ | 0.048 |
| **Section** | | | $\mathbf{F_{8,824.89}=32.20}$ | **<0.001** |
| **Year** | **-0.395** | **0.065** | $\mathbf{F_{1,46.29}=36.86}$ | **<0.001** |
| | | 700 m. buffer | | |
| Nº blue tits | 0.036 | 0.019 | $F_{1,2584.78}=3.416$ | 0.064 |
| **Section** | | | $\mathbf{F_{8,823.75}=32.27}$ | **<0.001** |
| **Year** | **-0.396** | **0.065** | $\mathbf{F_{1,46.54}=36.90}$ | **<0.011** |
| | | 800 m. buffer | | |
| Nº blue tits | 0.030 | 0.021 | $F_{1,2812.39}=2.087$ | 0.148 |
| **Section** | | | $\mathbf{F_{8,845.93}=30.32}$ | **<0.001** |
| **Year** | **-0.394** | **0.065** | $\mathbf{F_{1,46.87}=36.54}$ | **<0.001** |
| | | 900 m. buffer | | |
| Nº blue tits | -0.034 | 0.022 | $F_{1,3220.1}=2.299$ | 0.129 |
| **Section** | | | $\mathbf{F_{8,860.3}=29.60}$ | **<0.001** |
| **Year** | **-0.397** | **0.065** | $\mathbf{F_{1,47.3}=36.73}$ | **<0.001** |
| | | 1000 m. buffer | | |
| Nº blue tits | 0.038 | 0.024 | $F_{1,3683.8}=2.499$ | 0.113 |
| **Section** | | | $\mathbf{F_{8,878.7}=29.27}$ | **<0.001** |
| **Year** | **-0.399** | **0.065** | $\mathbf{F_{1,47.7}=36.96}$ | **<0.001** |
| | | 1250 m. buffer | | |
| Nº blue tits | -0.021 | 0.028 | $F_{1,3527.5}=0.615$ | 0.433 |
| **Section** | | | $\mathbf{F_{8,869.5}=31.05}$ | **<0.001** |
| **Year** | **-0.372** | **0.065** | $\mathbf{F_{1,48.7}=32.17}$ | **<0.001** |
| | | 1500 m. buffer | | |
| Nº blue tits | -0.046 | 0.027 | $F_{1,2165.40}=2.727$ | 0.098 |

| | | | | |
|---|---|---|---|---|
| **Section** | | | **$F_{8,838.21}$=32.40** | **<0.001** |
| **Year** | **-0.362** | **0.065** | **$F_{1,48.62}$=30.43** | **<0.001** |
| 1750 m. buffer | | | | |
| Nº blue tits | -0.049 | 0.029 | $F_{1,1662.38}$=2.958 | 0.085 |
| **Section** | | | **$F_{8,831.52}$=32.67** | **<0.001** |
| **Year** | **-0.360** | **0.065** | **$F_{1,48.84}$=30.102** | **<0.011** |
| 2000 m. buffer | | | | |
| Nº blue tits | -0.057 | 0.032 | $F_{1, 1688.89}$=3.050 | 0.080 |
| **Section** | | | **$F_{8, 853.06}$=32.68** | **<0.001** |
| **Year** | **-0.355** | **0.066** | **$F_{1,50.19}$=28.87** | **<0.001** |
| 2500 m. buffer | | | | |
| Nº blue tits | -0.051 | 0.042 | $F_{1, 1061.66}$=1.455 | 0.227 |
| **Section** | | | **$F_{8,876.82}$=32.00** | **<0.001** |
| **Year** | **-0.354** | **0.068** | **$F_{1,55.58}$=26.61** | **<0.001** |
| 3000 m. buffer | | | | |
| **Nº blue tits** | **-0.142** | **0.050** | **$F_{1,419.43}$=7.876** | **0.005** |
| **Section** | | | **$F_{8, 843.79}$=32.839** | **<0.001** |
| **Year** | **-0.293** | **0.072** | **$F_{1,59.12}$=16.15** | **<0.001** |
| 4000 m. buffer | | | | |
| Nº blue tits | -0.033 | 0.086 | $F_{1,44.28}$=0.149 | 0.701 |
| **Section** | | | **$F_{8,817.52}$=31.92** | **<0.001** |
| **Year** | **-0.360** | **0.087** | **$F_{1,44.27}$=17.01** | **<0.001** |

### SM-7: Association between breeding density during the nestling stage and adult mass

| | Estimate | SE | *F* | *P* |
|---|---|---|---|---|
| | Adult body mass (n=9306) | | | |
| **Great and blue tit density (3000 m.)** | **-0.034** | **0.015** | $F_{1,4246.8}$=4.926 | **0.026** |
| **Section** | | | $F_{8,6873.7}$=7.387 | **<0.001** |
| **Year** | **-0.203** | **0.024** | $F_{1,85.8}$=69.01 | **<0.001** |
| **$Age_{juv}$** | **-0.273** | **0.015** | $F_{1,6526.3}$=303.39 | **<0.001** |
| **$Sex_{male}$** | **-0.327** | **0.015** | $F_{1,5819.4}$=248.7 | **<0.001** |

### SM-8: Phenotypic rates of trait change across taxa

We used PROCEED (v. 6.1), a dataset containing information of phenotypic rates of change across trait and taxa (Gorné, L.D. *et al.* 2025), to compare the trends found for adult and nestling great tit mass with others available in the literature.

First, we used the data and code provided in PROCEED to compute the phenotypic rates of changes in Haldanes for the data provided. Second, we used the computed rates of change to obtain several descriptive metrics of the phenotypic rates of change available in the literature not including any experimental studies or those coming from the comparison of two different populations (i.e. synchronic; SM-7 Table 1, Fig. 1 & 2)

SM-8 Table 1. Descriptive statistics of the Haldanes available at PROCEED v. 6.1. considering different groups and traits.

| Taxa | Traits | Median | Abs. Median | Abs. Mean | Min. | Max. | n |
|---|---|---|---|---|---|---|---|
| Vertebrates | All | -0.002 | 0.070 | 0.304 | -27.20 | 6.721 | 2365 |
| Birds | All | -0.001 | 0.007 | 0.063 | -1.528 | 0.595 | 331 |
| Birds | Morphological | -0.001 | 0.007 | 0.073 | -1.528 | 0.595 | 231 |
| Bird | Mass | -0.055 | 0.063 | 0.271 | -1.528 | 0.063 | 13 |

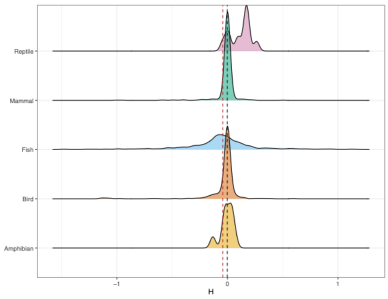


SM-8 Fig. 1. Distribution of the estimated Haldanes for vertebrates available at PROCEED v. 6.1. Extreme values (outside the central 95% of the distribution) were excluded for visualization purposes.

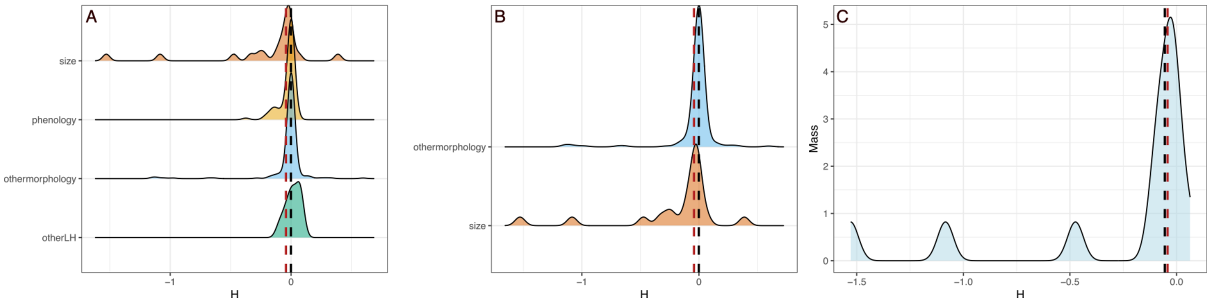


SM-8 Fig. 2. Distributions of the estimated Haldanes available at PROCEED v. 6.1. A) all Haldanes from birds split in four categories, B) Haldanes from morphological traits and C) Haldanes for mass related traits. The red dashed line represents the rate of change in body mass of great tits in Wytham woods, the black dashed line represents the median rate of change in each category
